# Suppression of insulin stimulated muscle glucose uptake by non-caloric sweetener sucralose and its reversal by an antidiabetic nutraceutical supplement

**DOI:** 10.1101/2022.11.17.516874

**Authors:** R.N Arpitha Reddy, K Saranya, Sanjana Battula, Gopi Kadiyala, Subramanian Iyer, Subrahmanyam Vangala, Satish Chandran, Uday Saxena

**Affiliations:** Reagene Innovations Pvt Ltd, Hyderabad, India; Kyntox Bio, Bangalore, India; Prodigy Bio, Florida, USA

## Abstract

Non caloric sweeteners (NCS) have been used for decades now as sugar substitutes in foods and beverages. The market for such products has grown immensely over time. There are human studies which report their negative effects on glucose metabolism with various results of disturbances in glucose metabolism, weight gain etc.

No studies to the best of our knowledge have directly addressed the impact of the NCS on muscle glucose uptake. Muscle tissue can account for over 70 percent of whole-body glucose uptake. Therefore, we examined directly the effect of NCS on muscle cell glucose uptake. We find that aspartame moderately increased insulin stimulated glucose uptake by muscle cells in vitro. But sucralose, saccharin and stevia suppressed insulin stimulated glucose uptake.

Sucralose is one of the most often used sweetener in foods and beverages globally it is important to understand its effects on glucose metabolism. Therefore, we explored the mechanism of its inhibition of glucose uptake by using an anti-diabetic nutraceutical which is known to target insulin mediated glucose uptake and metabolism pathways. We show here that the nutraceutical is able to relieve the suppression by sucrose in muscle uptake by a novel mechanism of action. We propose that such nutraceuticals may be useful to combine with sucralose containing products to offset negative effects of the NCS on glucose metabolism.

## Introduction

The use of NCS has increased dramatically over the last few years. With the incidence of pre-diabetes, type 2 diabetes and obesity globally, NCS are attractive candidates for substitution of sugar. This has encouraged their use in diet beverages, food and sweets. Millions of people consume NCS every day (1–4).

The effects of NCS in humans has been the subject of interest and studies report varied effects from lack of effects to negative impact on glucose metabolism (5–15). Most recently a clinical study has shown that aspartame and stevia had no impact on whole body glucose metabolism but sucralose and saccharin were shown to suppress glucose disposal (1). The mechanism of these effects was attributed to their effect on gut microbiome and dysbiosis.

To gain insights into the effects of NCS on glucose metabolism, we used an in vitro muscle cell system to study impact of the sweeteners on glucose uptake and metabolism. We did this since it is easier to approach mechanisms in a less complex system such as muscle directly as well as because muscle dominates in glucose uptake and metabolism in humans.

We find that both sucralose, saccharin and stevia and but not aspartame suppress insulin stimulated muscle cell glucose uptake. Furthermore, the effects of sucralose can by improved addition of an anti-diabetic nutraceutical by novel mechanisms(s).

## Methods

### MATERIALS AND METHODS

Glucose uptake studies using L6 cells were performed similar to our previous reports. The concentrations of Vitamin D3, Niacinamide and Lipoic acid were typically similar to the Required Daily Allowance (RDA) for each.

Briefly, L6 cells (NCCS) were maintained in Dulbecco’s modified Eagle’s medium (DMEM) supplemented with 10% Fetal Bovine Serum (FBS) and 50U Penicillin and 50μg Streptomycin at 37°C in an atmosphere of 5% CO_2_.

#### 2-NBDG Glucose uptake

2-(N-(7-Nitrobenz-2-oxa-1,3-diazol-4-yl)amino)-2-deoxyglucose [2-NBDG][Molecular probes - Invitrogen, CA, USA] was used to assess glucose uptake in L6 cells. Cells were seeded at 1X10^4^ density in 96 well plate and incubated at 37°C in an atmosphere of 5% CO_2_. After 24hours, cells were treated with Non-caloric sweeteners, Divitiz, Vitamin D, Niacinamide and Lipoic acid and kept for overnight incubation. Once after the incubation, cells were kept in Glucose-free medium for an hour before insulin stimulation. Cells were stimulated with 40IU insulin for 0, 10 and 20min and then incubated with 100μM of 2-NBDG for 20 minutes. The reaction was stopped by washing with cold 1X PBS and the cells were lysed using 0.1% Triton X-100. The lysate was then used to read the fluorescence intensity at an excitation of 485nm and emission of 535nm. Data presented represent an average of 3 to 6 separate experiments.

## Results

### 1. Effects of various non caloric sweeteners on insulin stimulated muscle glucose uptake

Shown in Figure 1 are the effects of various NCS (1 mM) on muscle glucose uptake. Aspartame did not have any decreasing effect and in fact showed an increase in glucose, 102% and 199% increase at 10 and 20 minutes after insulin stimulation. Sucralose demonstrated reduced uptake at 10 minutes after insulin stimulation (no increase relative to time zero) and at 20 minutes there was increase but only 112%. Saccharin was even worse than sucralose, showing complete block at 10 and 20 minutes after insulin stimulation. Uptake of glucose in presence of Stevia relative to time zero of insulin stimulation showed no improvement as well. For reference, under control condition (no NCS added), the increase in glucose uptake is 108% and 124% at 10 minute and 20 minutes respectively. These data suggest that saccharin, sucralose and stevia in that order suppress glucose uptake but aspartame does not suppress insulin stimulated uptake by muscle cells.

**Figure 1A:**
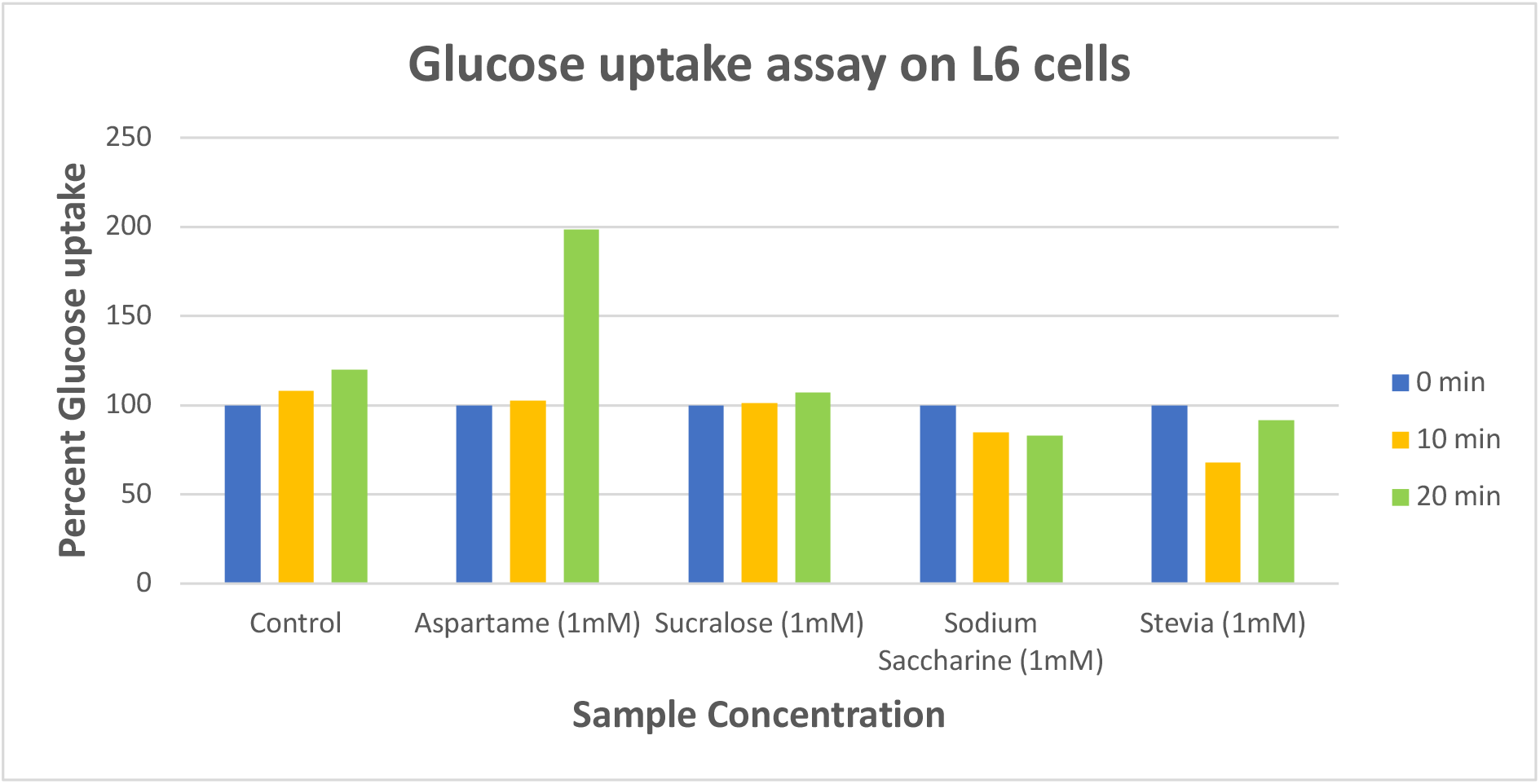
Effect of NCS on insulin induced glucose uptake. Data is shown as percent of control (time 0) at 0, 10 and 20 minutes after insulin stimulation

### 2. Alleviation of sucralose induced suppression of glucose uptake by an antidiabetic nutraceutical supplement

Since sucralose is the most used NCS we focused our attention in understanding the mechanism of its action in inhibiting insulin stimulated glucose uptake. We did not pursue saccharin since it is not in regular use due to potential carcinogenicity and stevia is a plant derived product which is composed of multiple chemical entities.

We have recently shown that a defined nutraceutical composition (called Divitiz) is able to improve insulin stimulated glucose uptake in muscle cells and overall glucose disposal in vivo (17). The mechanism of increase by the nutraceutical was due to targeted improvements in glucose uptake, glucose cellular metabolism and redox stress. We used this nutraceutical as a tool to explore its potential to improve upon glues uptake in presence of sucralose. Figure 2 A shows a typical effect of the nutraceutical on insulin stimulated glucose uptake.

**Figure 2A:**
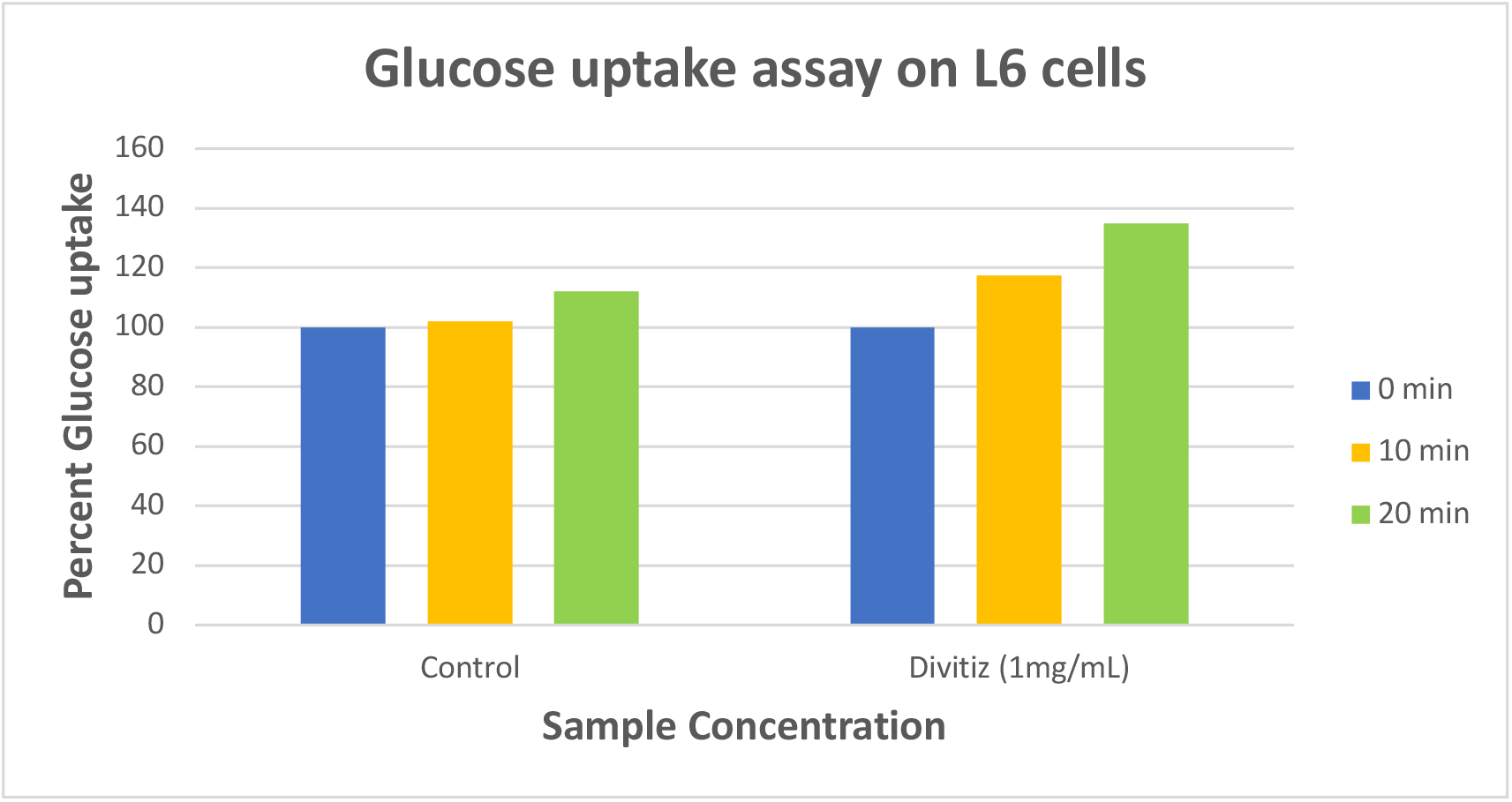
Effect of nutraceutical (Divitiz) on insulin induced glucose uptake relative to no nutraceutical added. Data is shown as percent of control (time 0) at 0, 10 and 20 minutes after insulin stimulation

With regards to NCS, as shown in Figure 2 B it was found that sucralose (100 uM) alone showed increase of 103% at 10 minutes and no significant increase at 20 minutes. When the nutraceutical supplement was added together with sucralose, the supplement increased insulin stimulated uptake by **152% at 10 minutes and 254% percent at 20 minutes** in presence of sucralose.

**Figure 2B:**
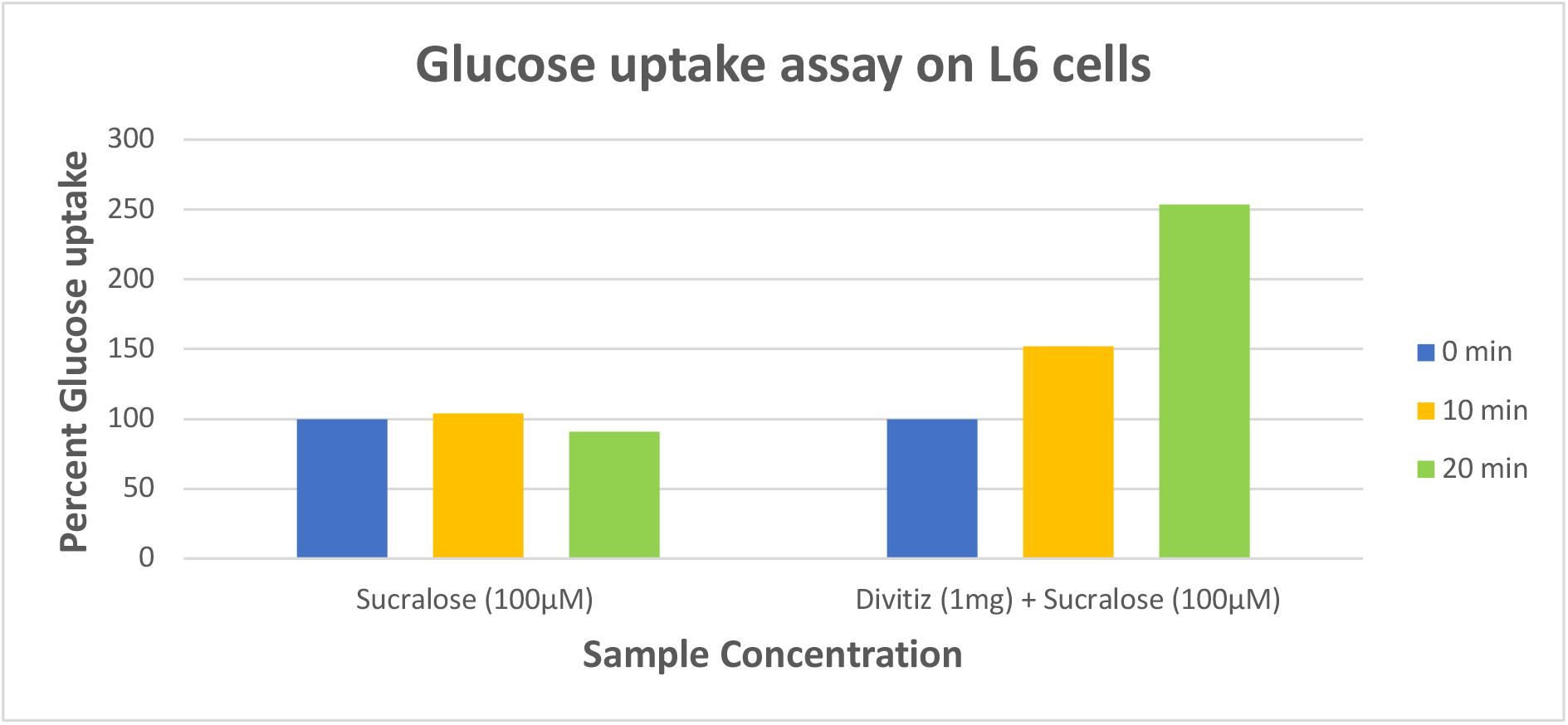
Effect of nutraceutical (Divitiz) on insulin induced glucose uptake in the presence of sucralose. Data is shown as percent of control (time 0) at 0, 10 and 20 minutes after insulin stimulation

We also explored the effects of sucralose at higher concentration (1 mM) and found that even high sucralose alone suppressed insulin stimulated at 10 minutes and the increase at 20 minutes was only 112%. Addition of the nutraceutical improved glucose uptake by **119 % at 10 minutes and 122%.** These data suggest that the sucralose induced suppression of glucose uptake can be overcome by using a nutraceutical which works by improving muscle glucose metabolism.

**Figure 2C:**
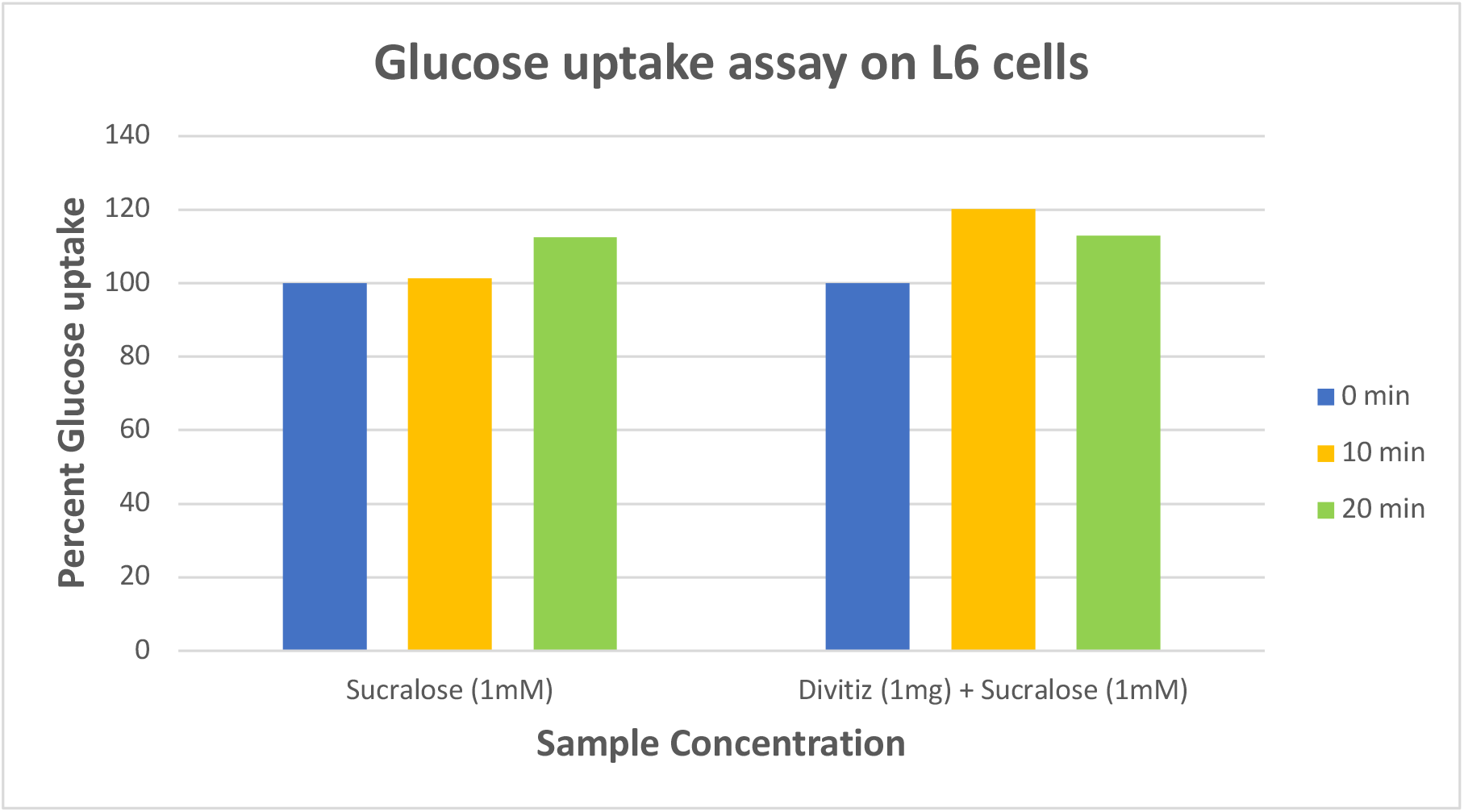
Effect of NCS (Divitiz) on insulin induced glucose uptake in the presence of high sucralose. Data is shown as percent of control (time 0) at 0, 10 and 20 minutes after insulin stimulation

### 3. Nutraceutical ingredients selectively improve sucralose induced suppression of glucose uptake

The nutraceutical we used here is composed of three ingredients, Vitamin D3 to improve glucose transporter GLUT4 levels thus increasing glucose uptake into the cells, Niacinamide which improves cellular tricarboxylic acid to generate energy from glucose and Lipoic acid, an antioxidant which decreases mitochondrial stress and improves glucose utilization by cells.

Since the nutraceutical as a composite was able to relive the suppressive effects, we used its ingredients to tease out where specifically sucralose may be causing the block in glucose uptake. Each of the ingredients was added in presence of sucralose to the cells and then insulin stimulated glucose uptake was measured. As shown in Figure 3, the insulin stimulated increase in presence of sucralose plus Vitamin D3 increased by 102% at 10 minutes but more robustly by **152% at 20 minutes**. Similarly, the addition of lipoic acid to sucralose increased the uptake by **142% at 20** minutes but no increase at 10 minutes after insulin stimulation. Interestingly niacinamide could not overcome the suppression induced by of sucralose with changes of 108% and zero percent at 10 and 20 minutes respectively. Collectively these data suggest that sucralose could be blocking glucose uptake at the GLUT4 transporter level as well as may be creating mitochondrial oxidative stress to suppress glucose metabolism. These data are insightful in deciphering the potential mechanism by which sucralose suppresses insulin stimulated muscle cell glucose uptake.

**Figure 3:**
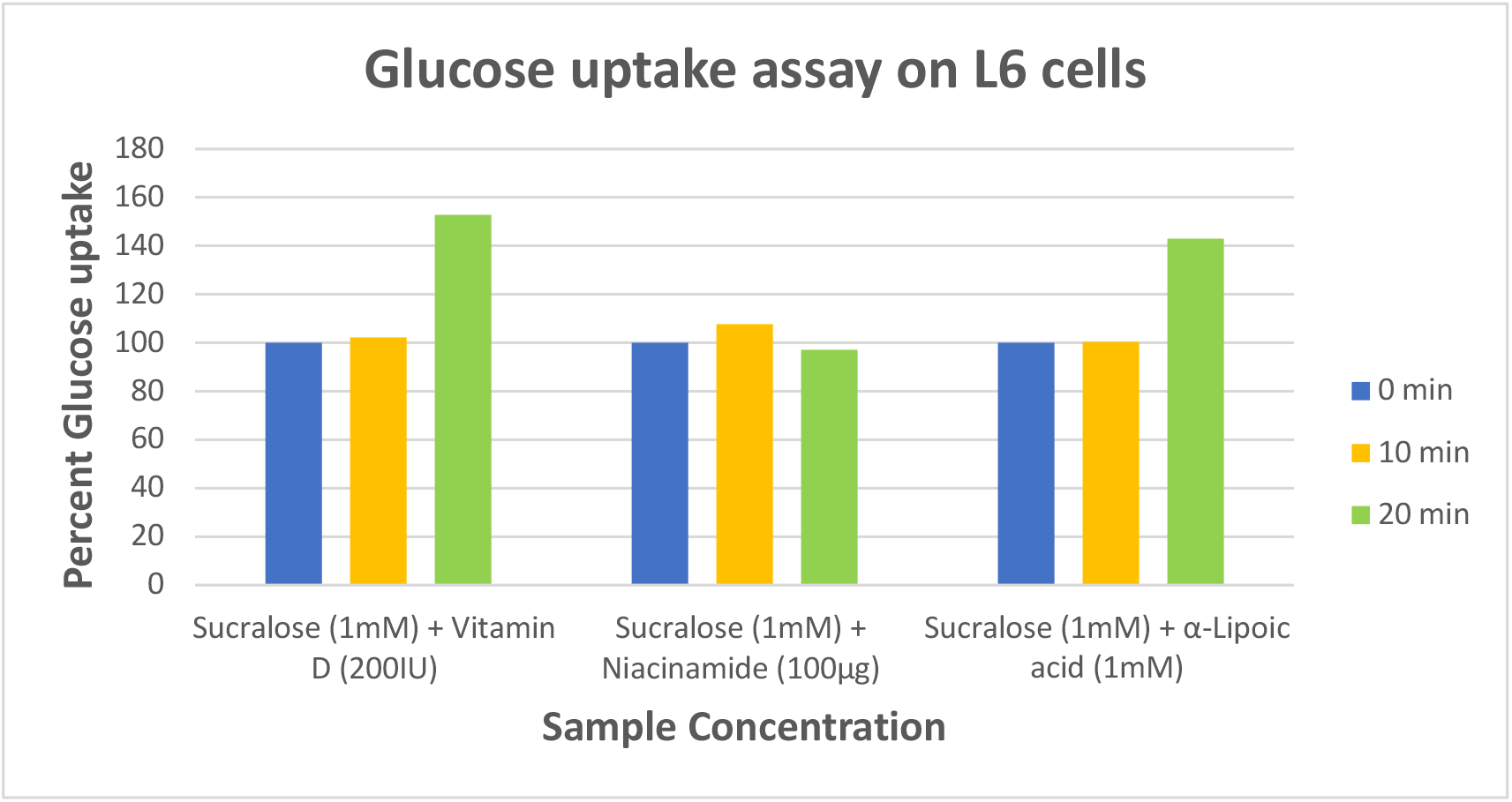
Effect of NCS individual ingredients on insulin induced glucose uptake in the presence of high sucralose. Data is shown as percent of control (time 0) at 0, 10 and 20 minutes after insulin stimulation

## Discussion

Our data presented here show that sweeteners sucralose, saccharin and stevia can potentially block insulin stimulated muscle uptake, which could ultimately result in hyperglycemia upon chronic usage. In this respect there is a recent clinical study which suggests that sucralose and saccharin decrease whole body glucose disposal, the same study showed that stevia and aspartame did not change glucose metabolism (1). The suggested changes were proposed to be due to effects of these sweeteners on gut microbiome and dysbiosis. Our in vitro data corroborate those negative clinical effects for sucralose and saccharin and lack of suppressive effect of aspartame.

We became interested in understanding the mechanism of suppression by sucralose for two reasons – one, it is commercially one of the most frequently used sweeteners and secondly the structure of sucralose is very similar to naturally occurring sucrose sugar with the only change being in one halogen atom. We hypothesized that such a small change may allow it to interfere with glucose uptake or muscle cell metabolism. Indeed, we found that the nutraceutical components Vitamin D3 which works by increasing GLUT4 transporter and lipoic acid which blocks redox stress in mitochondria were effective in relieving the effects of sucralose.

In our system, the suppression by sucralose may seem modest but if the sweetener is consumed chronically then the cumulative effects may eventually to lead to severe disturbance in glucose metabolism and ultimately hyperglycemia. Our data would suggest that inclusion of the antidiabetic nutraceutical may offset some of the effects of sucralose.

